# Evaluating the use of siRNA to silence the expression of the H5N2 virus polymerase genes as strategy to block the transmission of the avian H5N2 virus in mammalian cells

**DOI:** 10.64898/2026.05.04.722578

**Authors:** Richard Sutejo, Boon Huan Tan, Richard J Sugrue

## Abstract

We prepared siRNA libraries against the H5N2 virus NP gene, and the PA, PB1 and PB2 genes that express the proteins that form the virus polymerase complex. The antiviral activity of the siRNA libraries in H5N2 virus infected cells was initially assessed by using qPCR to measure the corresponding mRNA levels in the siRNA-treated cells. In this way siRNA molecules within each library were identified that exhibited to a greater than 70% reduction in levels of each target mRNA. A selection of these siRNA molecules was further evaluated for their antiviral activity in a multi-cycle H5N2 MDCK cell model. The siRNA molecules identified were successful in blocking virus transmission and lead to a reduction in influenza virus progeny virus production. This antiviral activity correlated with both the inhibition of nuclear export of the newly formed RNP complex’s that arise from the transcriptional activity of the input virus, and the inhibition of the polymerase activity of the newly formed virus polymerase complexes. This study highlights the potential use of siRNA as a strategy to block virus transmission by targeting the avian influenza virus polymerase complex.

## Introduction

Avian influenza viruses (AIV) are maintained in the feral aquatic bird populations and these animals are thought to be the reservoir for the influenza A viruses that infect other avian and non-avian hosts [1]. Infection of domestic poultry by AIV is of major economic importance, and in this context infection of poultry by highly pathogenic avian influenza (HPAI) viruses (e.g. H5N1) can lead to high fatality rates in poultry flocks. However, AIV can also infect non-avian hosts, including humans [2, 3, 4], and while HPAI infection in humans is often associated with high mortality rates, associated fatalities due to human transmission of low pathogenic avian influenza (LPAI) viruses (e.g. H5N2 virus) are rare. However, virus transmission and virus pathology are separate facets of the AIV and human interaction, and transmission of AIV to humans does not necessarily lead to high mortality rates. Poultry workers have tested seropositive for avian H5N2 and H9 viruses suggesting prior infection [5, 6], and H9N2 infection in humans is usually associated with mild influenza-like-illness (e.g. [2]). Therefore, while transmission of AIV to humans is possible, the capacity of these viruses to cause high fatality rates in humans is not shared by all AIVs. However, in all cases the AIV need to be able to adapt to their new host to enable transmission to the new host, and a major focus of research is geared towards understanding the process of species adaptation in the context of facilitating virus transmission in different hosts.

The influenza A virus genomic RNA (vRNA) contains eight RNA segments that are single-stranded and of negative polarity, and these eight segments give rise to all the virus structural and non-structural proteins. The virus polymerase (pol) complex is a multi-protein complex consisting of three virus-expressed polymerase (P) proteins, namely the PA, PB1 and PB2 polymerase proteins. The pol complex synthesizes new copies of vRNA and produces short methylated capped RNA primers from cellular mRNA that are used to prime virus mRNA synthesis by the process of cap-snatching. Distinct biochemical properties are associated with each P protein and the fully functional pol complex requires all the activities associated with the individual P proteins. The PB2 protein contains a cap-binding domain that immobilizes the host-cell mRNA at the replication complex, while endonuclease activity within the PA protein generates the methylated capped primers that are used in virus gene transcription. The PB1 protein contains the polymerase activity, the 5′ and 3′ binding sites for the vRNA, and the interaction domains for the PA and PB2 proteins. The structural analysis of the pol complex has provided mechanistic insights on how these different activities are orchestrated [7, 8, 9,10,11]. Although the capacity of influenza virus to infect different hosts is dependent upon several factors, the biological properties of the individual virus protein that form the pol complex is a major factor in determining host range i.e. the maintenance of the virus in a specific host [12, 13]. Specific amino acid sequence motifs within the different proteins that form the RNP complex have been identified that are associated with host adaptation. While the biological mechanisms that are underpinned by these specific sequence motifs during the host adaptation have been proposed in some specific cases [14, 15], in general these mechanisms remain poorly defined.

The virus nucleoprotein (NP) coats the single-stranded virus genomic RNA of negative polarity (vRNA), which together with the pol complex forms a larger independent transcriptional unit called the ribonucleoprotein (RNP) complex [16]. In virus-infected cells the newly formed RNP complexes are formed in the nucleus, and these are exported from the nucleus through the nuclear pore complex via the exportin 1/chromosome region maintenance 1 (crm1) export pathway. The RNP complex also associates with the virus expressed nuclear export protein (NEP), and it is the interaction between the NEP and the nuclear pore complex that mediates the nuclear export of the RNP complexes into the cytoplasm [17]. The individual RNP complexes representing each of the eight virus gene segments are subsequently exported to the cell surface at the site of virus particle assembly. We have also previously noted nuclear retention of the newly formed RNP complexes in H1N1/WSN virus-infected cells when siRNA was used to artificially silence the expression of the PA protein [18]. This indicated the requirement for efficient PA protein expression to enable the efficient nuclear export of the newly formed RNP and suggested that equimolar expression of the virus P proteins may be a requirement for efficient nuclear export of the newly formed RNP complexes. This also suggested that silencing the expression of the polymerase genes may be a viable antivirus strategy to block virus transmission.

The avian H5N2 virus (A/duck/Malaysia/F118/2004) was isolated from live poultry during routine a surveillance program [19]. The H5N2 virus could be propagated in both the avian and mammalian derived cell types (e.g. MDCK cells) [20], which suggested that although the H5N2 virus was of avian origin, it had adapted to replicate in a non-avian cell environment. In this current study we have therefore examined the effects of down regulating the expression of the P proteins on the propagation of the H5N2 virus in MDCK cells by using a selective and targeted siRNA approach to artificially silence P gene expression. This study allowed us to evaluate the use of siRNA molecules as an antiviral strategy to target the virus polymerase protein and prevent transmission of the avian H5N2 virus in non-avian cells.

## Methods

1. **Cells and viruses used.** The MDCK and HEK 293T cells were maintained in DMEM (Gibco) supplemented with 10% FCS (Gibco) and penicillin and streptomycin (pen/strep) (Gibco) at 37°C in a humidified chamber at 5%CO_2_. Chick embryo fibroblasts (CEF) were prepared from 8 to 10-day-old chick embryos and maintained in DMEM containing 10% FBS and pen/strep as described previously [20]. The isolation of the H5N2 (A/duck/Malaysia/F118/2004) was described previously [19] and the H1N1/WSN was purchased (ATCC). Both viruses were propagated in 7 day old embryonated eggs and the infected allantoic fluid harvested. Aliquots of the virus stocks were stored at -80°C and aliquots thawed prior to use. All work using the H5N2 (A/duck/Malaysia/F118/2004) virus was carried under BSL3 laboratory conditions in compliance with local regulations.
2. **Reagents.** The anti-NP antibody (Chemicon), anti-Lamin A/C antibody (Santa Cruz Biotechnology) and Evans Blue cell stain (Sigma) were purchased. The anti-mouse and anti-rabbit IgG conjugated to Alexa Fluor™ 488 and Alexa Fluor™ 555 were also purchased (ThermoFisher). Preparation of the antibodies against the influenza virus PA, PB1 and PB2 proteins were described previously [21].
3. **siRNAs and transfections.** All siRNAs were synthesized by Dharmacon (USA). The H5N2 specific siRNAs were produced as SMARTpool libraries that target the PA, PB1, PB2 and NP genes of the H5N2 virus (A/duck/Malaysia/F118/2004) [19], and the siGFP were synthesised using the sequence published previously [22]. The RNA sequences for all siRNA molecules are presented (Table. 1). The siRNA transfections were optimised using siGLO (Dharmacon) and transfections were performed with a 100nM concentration for each siRNA in Lipofectamine 2000.

### Virus infection

In the single cycle infections the cells were infected with each virus using a multiplicity of infection (moi) of 5 in DMEM (Gibco) supplemented with 2% (v/v) FCS (Gibco) and pen/strep (Gibco). Mock-infected cells and the infected cells were maintained in DMEM (Gibco) supplemented with 2% (v/v) FCS (Gibco) and pen/strep (Gibco). The multiple cycle infections were performed using a moi of between 0.1 and 0.01 as indicated, and the infection performed in DMEM supplemented with pen/strep, 0.21%(w/v) BSA, 1μg/ml trypsin-TPCK. In all cases the cells were incubated at 37°C in a humidified chamber at 5%CO_2_ until the time of harvesting.

### Plaque assay

Virus infectivity was assessed using standard plaque assays. Virus preparations were serially diluted in PBS and added to confluent MDCK monolayers. An overlay was used that consisted of 1% (w/v) low-melting point agarose (Invitrogen) in DMEM and 0.21%BSA, 1μg/ml trypsin-TPCK. Cells were incubated at 37°C in a humidified chamber at 5%CO_2_ until the time of appearance of the virus plaques in the MDCK cell monolayers.

### Immunofluorescence microscopy

The cells were fixed using either 4% (w/v) paraformaldehyde (PFA) in PBS and after washed using PBS at 4°C the PFA-fixed cells were permeabilised using 0.1% (v/v) triton X100 in PBS at 4°C for 15 mins prior to staining. The cells were stained using the appropriate primary and secondary antibody combinations (Alexa Fluor™ 488 or Alexa Fluor™ 555 as appropriate) and mounted on glass slides using mounting media (Dakocytomation Fluorescence Mounting Media, Dako, USA). The stained cells were examined using a Nikon Eclipse 80i Microscope (Nikon Corporation, Tokyo, Japan) with an Etiga 2000R camera (Q Imaging, Teledyne Photometrics, Tucson AZ, USA) attached. The images of immunofluorescence-stained cells were recorded using Q Capture Pro ver. 5.0.1.26 (Q Imaging, Teledyne Photometrics).

### Real-time quantitative PCR (qPCR)

The mRNA levels for each polymerase gene were determined as described previously [20] using the virus specific primers (Stable 1). The Real-time qPCR was performed using the LightCycler 2.0 System, software ver 5.32 (Roche) according to the previously published protocol [23]. The normalisation was performed using the primer pairs for the canine elongation factor.

### Minireplicon polymerase assay

This was performed as described previously using luciferase expression as the reporter assay [18]. The corresponding NP gene and polymerase genes for the H5N2/F118 virus was cloned into the pCAGGS mammalian vector using the virus-specific primers (STable. 2) to generate the vectors pCAGGS/H5N2-NP, pCAGGS/H5N2-PA, pCAGGS/H5N2-PB1 and pCAGGS/H5N2-PB2. The vector pPolL1-Luc carried the luciferase gene, and vector pPolL1-EGFP acted as a negative control in these assays. Transfections using Lipofectamine 2000 (Invitrogen) were performed in HEK 293T cells at 37°C and at 3 days post-transfection the tissue culture media was harvested and the luciferase activities measured. This was performed using a Gaussia luciferase assay kit and the measurements were made using a Fluoroscanskan Ascent FL (Thermo) reader.

## Results and Discussion

### 1. Detection of the nuclear export of the H5N2 virus RNP complex in virus infected cells by using imaging

This study was performed using the H5N2 (A/duck/Malaysia/F118/2004) avian virus isolate [18], and the MDCK cell line was used for the infections since the H5N2 virus has been shown to be permissive in this cell background [19]. The nuclear export of the RNP complex was primarily assessed using the anti-NP antibody to image virus-infected cells in the presence or absence of siRNA treatment. We confirmed that the MDCK cell line was fully permissive for infection with the H5N2 virus by comparing the replication kinetics of the H5N2 virus with the H1N1 virus using a multicycle infection (Fig 1). The MDCK cells were infected with each virus using a moi of 0.1 and 0.01 and the virus titers measured over 72 hrs. Although the H1N1 virus exhibited replication kinetics that were faster than the H5N2 virus in these cells, high levels of progeny virus was recorded for both viruses by 72hpi, albeit with an approximate 10-fold reduction in the H5N2 virus titer compared with the H1N1 virus (Fig. 1A). Similar virus titers were also achieved in both MDCK and CEF cells when these were infected with the H5N2 virus using a moi of 0.1 (Fig. 1B), although the virus replication kinetics appeared to be slightly faster in CEF cells which presumably reflects the avian origin of the H5N2 virus. Irrespective of these differences in recoverable virus infectivity, these confirmed that that the MDCK cell line was fully permissive for infection with the H5N2 virus.

**Figure 1.**
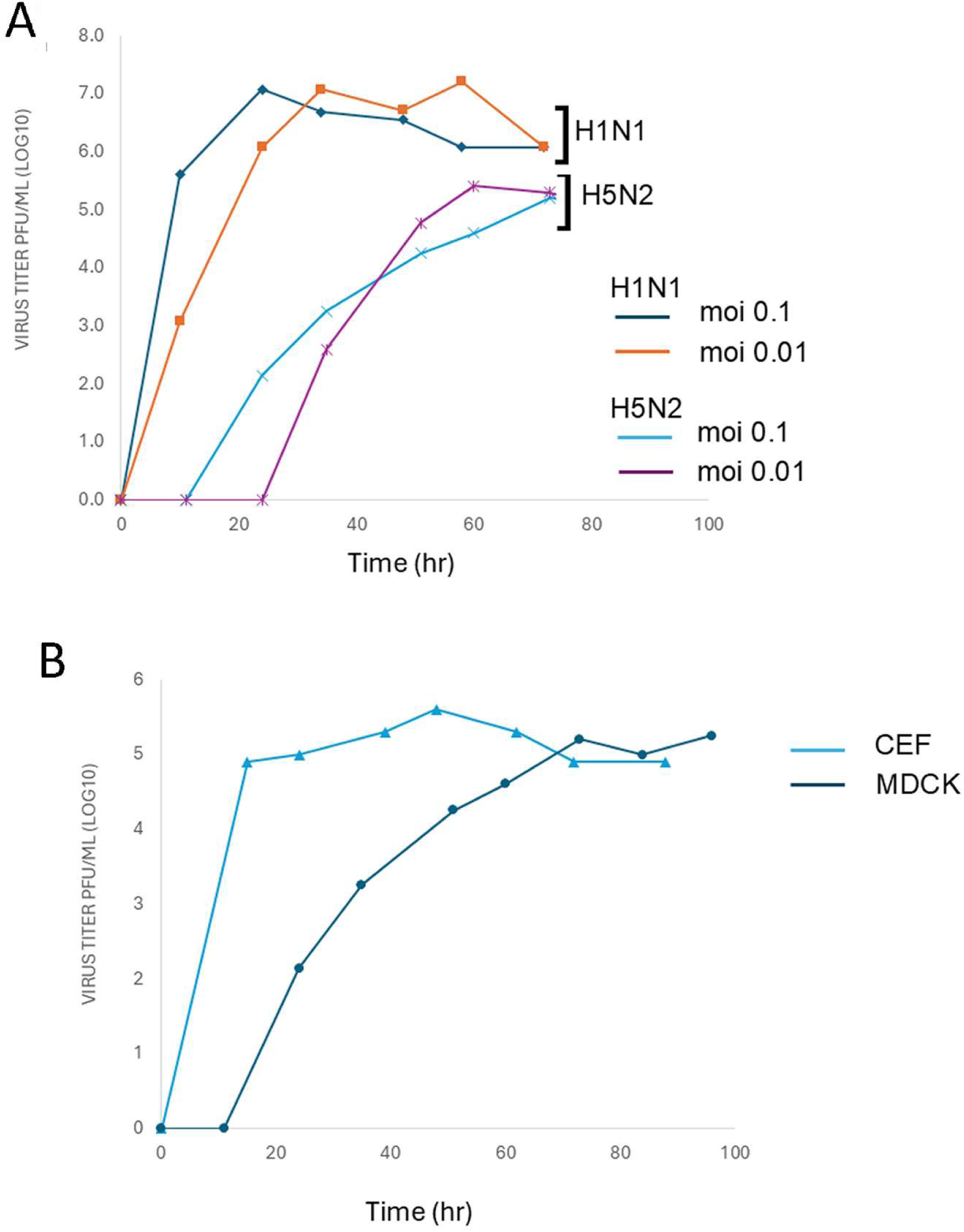
The MDCK cells are permissive for H5N2 virus infection. **(A)** MDCK cell monolayers were infected with the H1N1 and H5N2 viruses using the multiplicity of infection (moi) indicated. At each time point a portion of the virus supernatant was removed and the virus infectivity assessed using plaque assay. **(B)** MDCK and chick embryo fibroblasts (CEF) cells were infected with H5N2 viruses using a moi of 0.1. At each time point a portion of the virus supernatant was removed and the virus infectivity assessed using plaque assay. Each measurement is an average of three readings.

Cells were also either mock infected or infected with the H5N2 virus and at 24 hrs post-infection (hpi) the cells were co-stained using anti-NP and anti-Lamin A/C and examined by immunofluorescence (IF) microscopy (Fig. 2 A and B). The anti-Lamin A/C antibody detects the Lamin A and C isoforms that are present within the nuclear lamina, and this antibody therefore allows detection of the cell nucleus. The anti-NP staining in the H5N2 virus infected cells was distributed throughout the cell, both within the location demarcated by the anti-Lamin A/C-stained nucleus and within the cytoplasm (Fig. 2C), and this was consistent with nuclear export of the H5N2 virus RNP complexes in these cells. Cells were also infected with the H5N2 virus, and at between 8 and 16 hpi the cells were stained using anti-NP and the cell stain Evans Blue (EB) and imaged using IF microscopy (SFig. 1). EB is a non-specific cell stain that stains all the cells in the field of view i.e. both infected and non-infected, which allowed us to distinguish the non-infected and infected cells in the same field of view. This analysis allowed us to monitor the nuclear export of the RNP complexes in the virus-infected cells, from the anti-NP staining in the nucleus at the early stages of infection to the cytoplasmic anti-NP staining at the later stages of infection. While at the early stages of infection at 8 and 10 hpi the anti-NP staining pattern suggested a nuclear localization (SFig. 1B and C), by 16 hpi diffuse anti-NP staining across the whole cells was consistent with nuclear export of the RNP into the cytoplasm of the infected cells (SFig. 1D). These data collectively confirmed the compatibility of the MDCK cell line for H5N2 virus infection and indicated that anti-NP staining of H5N2 virus infected cells could be used to assess nuclear export of the newly formed H5N2 virus RNP complexes in virus infected cells following treatment with the different siRNA molecules.

**Figure 2.**
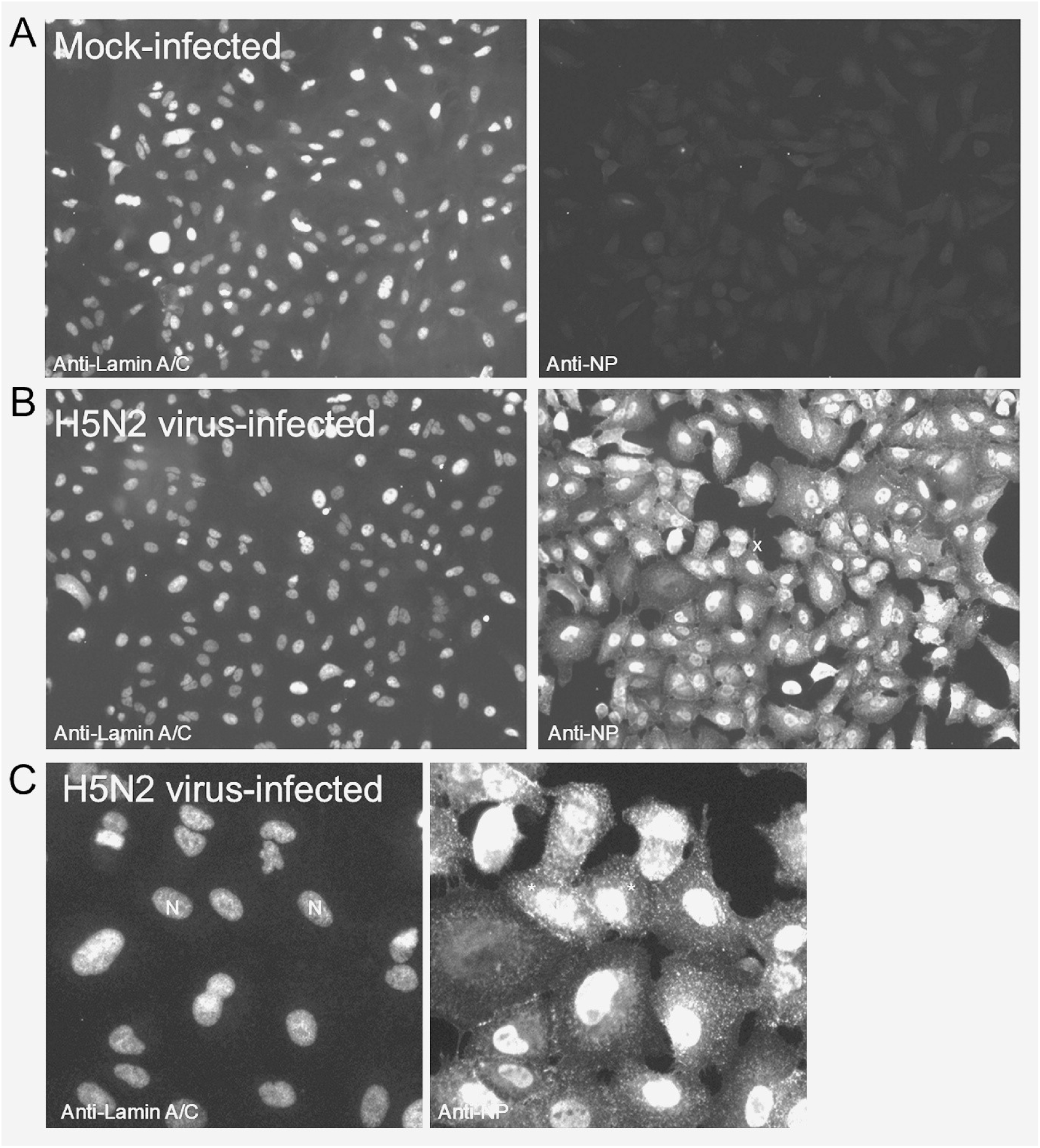
The RNP complex is exported into the cytoplasm in H5N2 virus-infected MDCK cells. MDCK cells were either **(A)** mock-infected or **(B)** H5N2 virus infected and after 24 hrs post-infection the cells were co-stained with anti-NP and anti-Lamin A/C and imaged using immunofluorescence microscopy (objective x 20 magnification). **(C)** is an enlarged image taken from the region in **(B)** indicated by x. In this image the anti-Lamin A/C-stained nucleus (N) and the corresponding cytoplasmic anti-NP staining (*) are highlighted.

### 2. The effects of silencing the expression of individual polymerase proteins on the nuclear export of newly formed RNP complexes in H5N2 virus-infected cells

A library of siRNA molecules was generated against the A/duck/Malaysia/F118/2004 H5N2 virus PB2, PB1, PA, and the NP genes. In each library a series of four siRNA molecules were designed using the sequences of segment-1 (siPB2 1-4), segment-2 (siPB1 1-4), segment-3 (siPA 1-4) and segment-5 (siNP 1-4) (Table 1A). The location of the individual siRNA molecules in each respective gene sequence are shown (Table 1B), and these were located across the length of each respective gene. The antiviral activity associated with each siRNA was primarily assessed in H5N2 virus-infected MDCK cells. Our previous study using siRNA suggested that the qPCR was a reliable method to examine the knock-down effect on virus gene expression following siRNA treatment [20]. These siRNA molecules were therefore initially evaluated for their ability to silence the expression of individual polymerase proteins by using qPCR to compare the mRNA levels in the siRNA-treated and untreated cells.

**Table 1.**
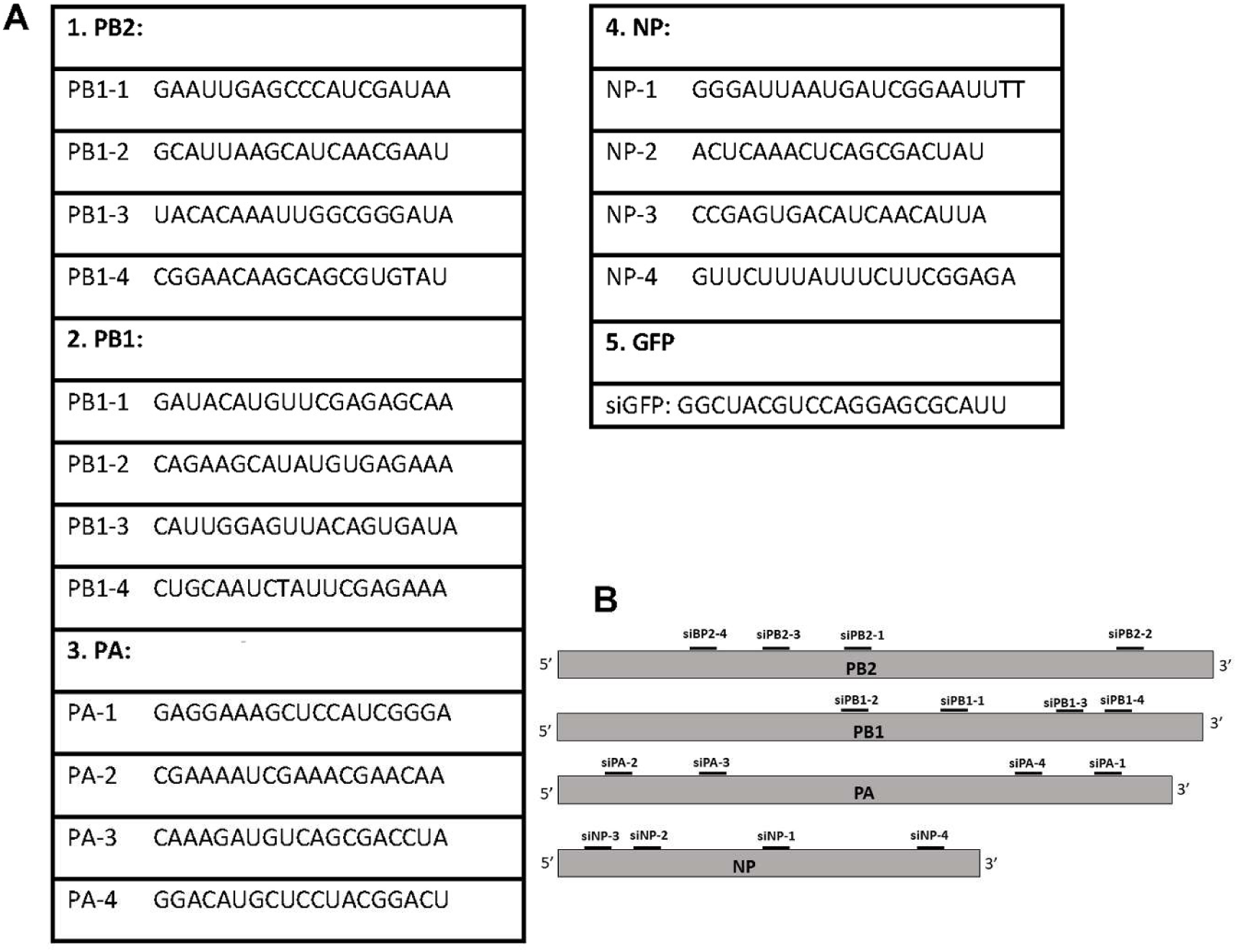
Sequences of the SMARTpool siRNA. (A) The sequence of the all the siRNA species used in this study. The SMARTpool for the PA, PB1, PB2 AND NP genes of the H5N2 virus were generated using the sequences of A/duck/Malaysia/F118/2004 [19]. **(B)** approximate location of the siRNA species that were present within the SMARTpool in the H5N2 virus coding region for each respective gene.

#### 2.2.1 Segment-5 (siNP-1 to -4)

MDCK cells were treated with siGFP and either siNP-1, siNP-2, siNP-3 or siNP-4 and infected with the H5N2 virus using a moi of 5, and at 24 hpi the levels of NP mRNA levels determined by qPCR and compared with the corresponding mRNA levels in non-treated cells (Fig. 3A). In siGFP-treated cells similar NP mRNA levels to that in non-treated cells were recorded. Although all siNP molecules showed reduced NP mRNA levels in the H5N2 virus infected cells, the siNP2 and siNP4 showed an approximate 30% reduction in NP mRNA levels, while the siNP-1 and siNP-3 treated cells showed by comparison an approximate 70% reduction in NP mRNA levels.

**Figure 3.**
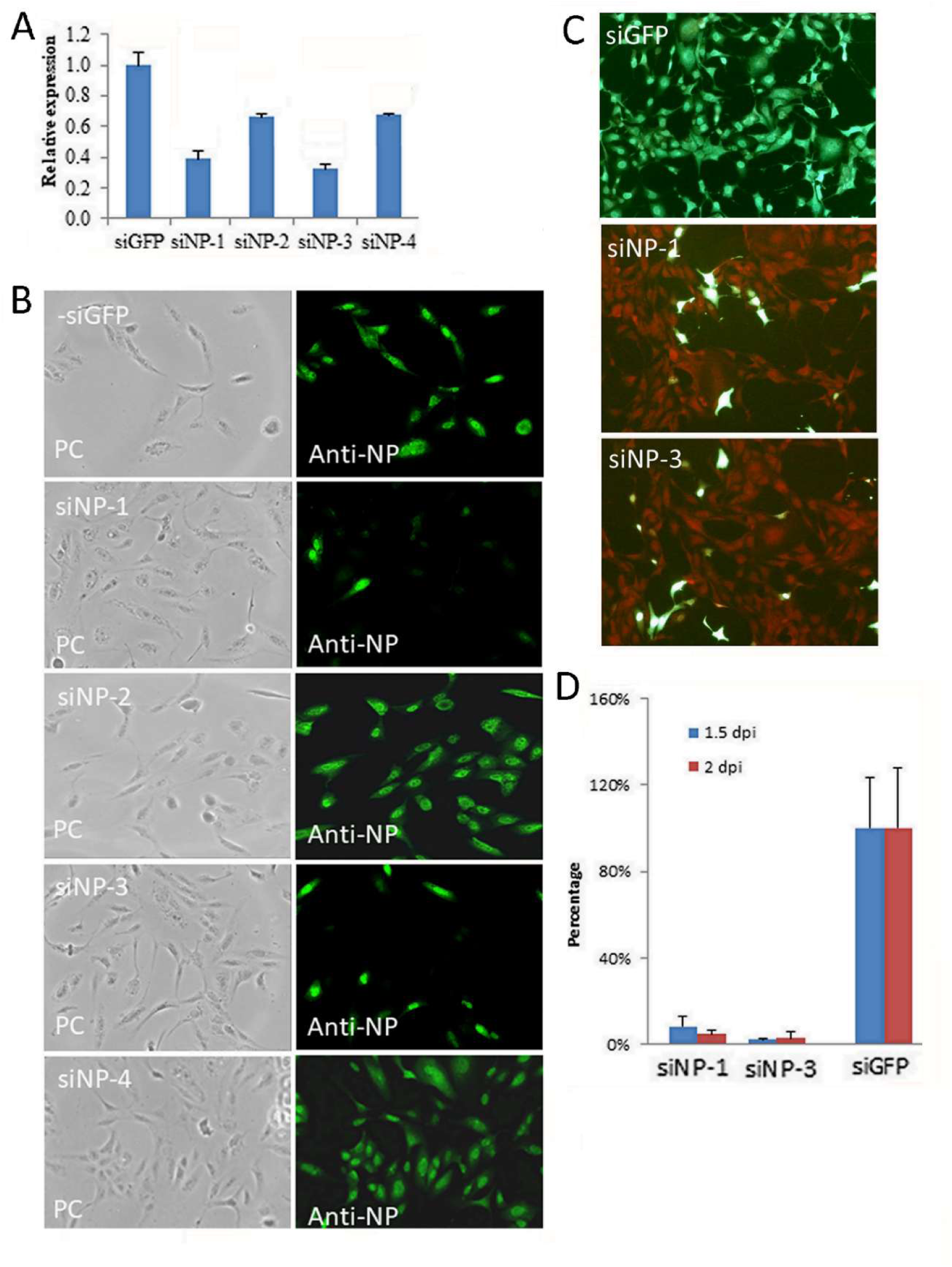
Silencing of the H5N2 virus NP protein in H5N2 virus-infected MDCK cells. **(A)** MDCK cells were treated with either siGFP or siNP-1, siNP-2, siNP-3 or siNP-4 as indicated and infected with the H5N2 virus using the multiplicity of infection of 5. The mRNA coding for the NP was measured by qPCR and the relative levels of each is shown with respect to the corresponding mRNA levels in untreated cells is shown. Each reading was performed in triplicate, and the average readings are shown. **(B)** MDCK cells were treated with either siGFP and each of the siNP species as indicated and infected with the H5N2 using the multiplicity of infection of 5. The cells were stained using anti-NP and imaged by fluorescent microscopy (objective x 40 magnification). Also shown is the same field of view imaged using phase contrast (PC) to enable imaging of all cells in the same field of view. **(C)**. MDCK cell monolayers were treated with siGFP, siNP-1 and siNP-3 and infected with the H5N2 virus using the multiplicity of infection of 0.1. After 48 hrs post-infection the cells were co-stained using anti-NP (green) and the cell stain Evans blue (red) and imaged using immunofluorescence microscopy (objective x 20 magnification). The Evans blue was used to visualise all the cells in the field of view, both the antibody-stained infected cells and the unstained and uninfected cells. **(D)** MDCK cell monolayers were treated with siGFP, siNP-1 and siNP-3 and infected with the H5N2 virus using the multiplicity of infection of 0.1. After 36 and 48 hrs post-infection the virus infectivity in the tissue culture medium was measured using plaque assay. The percentage change relative to that in untreated cells infected with the H5N2 virus is shown.

In a parallel analysis the cells treated with these siRNA molecules were infected with the H5N2 virus using a moi of 5 and after 24 hpi the cells were stained with anti-NP and imaged using phase contrast (PC) microscopy and IF microscopy (Fig. 3B). The PC microscopy allowed imaging of all cells in the field of view, while IF microscopy allowed imaging of the infected anti-NP-stained cells. While in siGFP-treated, siNP-2-treated and siNP-4-treated infected cells approximately 100% of the cells exhibited anti-NP-staining, in siNP-1-treated and siNP-3-treated cells no anti-NP staining was apparent in approximately 90-95% of the cells. This confirmed that the siNP-1-treatment and siNP-3 treatment inhibited the expression of the H5N2 virus NP gene expression in H5N2 virus-infected cells. Furthermore, NP associated with the RNP complexes in the input virus remained undetected in the siNP-1-treated and siNP-3-treated cells; presumably due to the NP levels arising from the input virus being below the levels of antibody detection. Interestingly, although an approximate 40% reduction in NP mRNA levels was noted in H5N2 virus-infected cells treated with siNP-2 and siNP-4, this relatively moderate reduction did not appear to correspond with an antiviral effect. This suggested that the magnitude of the reduction in NP mRNA levels is also an important factor in determining an antiviral effect for these siRNA molecules, although the molecular basis for this correlation is currently unclear. This was consistent with our previous analysis on the H1N1 virus when knock down of NP gene expression by siRNA treatment led to an 80-90% reduction in NP mRNA levels. This previous data also confirmed that qPCR could reliably be used to assess the silencing of virus gene expression following siRNA treatment.

The antiviral effects of the siNP-1 and siNP-3-treated cells were confirmed using a multiple cycle infection model in which cells were treated with siGFP and either siNP-1 or siNP-3 and infected with the H5N2 virus using a moi of 0.01. At 48 hpi the cell monolayers were co-stained using EB and anti-NP and the stained monolayers were imaged using IF microscopy (Fig. 3C). The siGFP-treated cells exhibited extensive anti-NP staining across the whole monolayer, which was consistent with efficient virus transmission in the cell monolayer, and cells exhibiting only exhibiting EB-staining were not apparent. In contrast, in the siNP-1 or siNP-3-treated cells reduced levels of anti-NP staining were observed, which was consistent with an inhibition of virus transmission. The low levels of anti-NP staining in the siNP-1-treated and siNP3-treated cells that we observed accounted for between 5 and 10% of the cells in the field of view. We presume that this is due to a minor population of cells that did not receive the siRNA molecule since under our experimental conditions the transfection efficiency of the siRNA molecule in these cells was not 100%. While these cells would be infectable with the H5N2 virus, subsequent transmission of the virus within the cell monolayers treated with siNP-1 and siNP-3 was blocked. In addition, the recoverable virus infectivity present in the tissue culture medium of H5N2 virus-infected cells treated with siGFP and either siNP-1 or siNP-3 was measured and compared with non-treated cells (Fig. 3D). No significant reduction in the recoverable virus infectivity in siGFP treated cells was noted, while in siNP-1-treated or siNP-3-treated cells there was an approximated 90% reduction in the levels of virus infectivity recovered. This residual infectivity detected in the siNP-1-treated or siNP-3-treated cells presumably reflects the presence of the minor population of cells in the cell monolayer that were not transfected with the siRNA molecules as discussed above.

#### 2.2.1 Segment-3 (siPA-1 to -4)

MDCK cells were treated with siGFP and either siPA-1, siPA-2, siPA-3 or siPA-4 and infected with H5N2 virus using a moi of 5, and at 24 hpi the PA mRNA levels were measured and compared with that in non-treated cells (Fig. 4A). Similar PA mRNA levels were detected in non-treated and siGFP-treated H5N2 virus-infected cells, which indicated no change in the PA mRNA levels following siGFP-treatment. Although all siPA molecules showed reduced PA mRNA levels, we noted differences in the magnitude of the changes in the expression levels. While siPA-2 showed an approximate 20-30% reduction in PA mRNA levels, cells treated with the siPA-1, siPA-3 and siPA-4 showed a 70-80% reduction in PA mRNA levels. In a parallel analysis cells treated with these siRNA molecules were infected with the H5N2 virus and at 24 hpi the cells were stained with anti-NP and imaged using IF microscopy (Fig. 4B). While anti-NP staining was observed in all cells infected with the H5N2 virus, in cells treated with siGFP and siPA-2 we noted a diffuse anti-NP staining pattern that was also observed in non-treated H5N2 virus-infected cells. This was consistent with the expression of the NP and the subsequent nuclear export of the newly formed RNP complexes from the nucleus of the infected cells. Although anti-NP staining was detected in cells treated with siPA-1, siPA-3 and siPA-4, the staining pattern was distinct from that observed in siGFP and siPA-2-treated cells. In H5N2 virus-infected cells treated with siPA-1, siPA-3 and siPA-4 in approximately 90-95% cells in the field of view the anti-NP staining appeared to be primarily localized to the nucleus of the infected cells. This staining pattern was identical to the staining patterns exhibited in the siPA-H1N1-treated H1N1 virus infected cells that were described previously [20]. The data obtained above using the siPA molecules was consistent with the synthesis of the newly formed anti-NP stained RNP complexes during H5N2 virus infection occurring via the transcriptional activity associated with the RNP complexes of the input virus. Furthermore, the anti-NP staining pattern indicated that the reduced expression of the PA protein following siRNA treatment was associated with impaired nuclear export of the newly formed H5N2 virus RNP complexes. This observation in the H5N2 virus-infected cells following silencing of the PA protein expression was similar to the effects of silencing the PA protein expression in the H1N1 virus-infected cells described previously [18] suggesting that these observations were not influenza virus type-specific.

**Figure 4.**
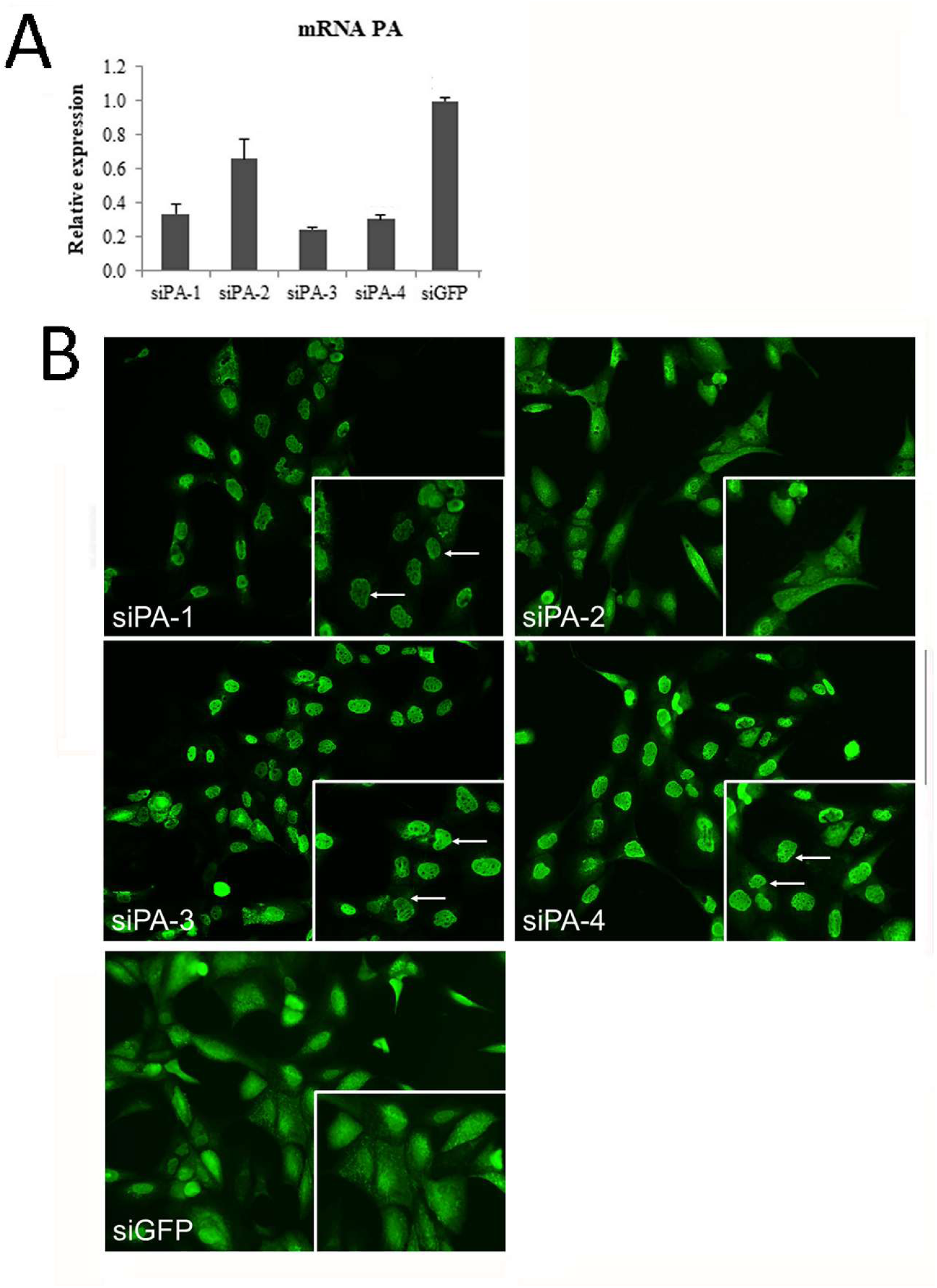
Silencing of the H5N2 virus PA protein in H5N2 virus-infected MDCK cells. **(A)** MDCK cells were treated with either siGFP or siPA-1, siPA-2, siPA-3 or siPA-4 as indicated and infected with the H5N2 virus using the multiplicity of infection of 5. **(A)** The mRNA coding for the PA was measured by qPCR and the relative levels of each is shown with respect to the corresponding mRNA levels in untreated cells is shown. Each reading was performed in triplicate, and the average readings are shown. **(B)** MDCK cells were treated with either siGFP and each of the siPA species as indicated and infected with the H5N2 using the multiplicity of infection of 5. The cells were stained using anti-NP and imaged by fluorescent microscopy (objective x 40 magnification). The insets in each plate are enlarged images of the cells in the field of view for each plate to more clearly image the anti-NP staining pattern and the nuclear anti-NP staining pattern (white arrows) is highlighted.

#### 2.2.3 Segment-1 and 2 (siPB1-1 to -4 and siPB2-1 to -4)

The data presented above indicated that the nuclear export of the newly formed RNP complexes required the expression of the PA protein, and the reason for this was not clear. While this may be related to the endonuclease activity in the PA protein, the RNP complex is a multiprotein complex with a defined stoichiometry of the different virus components involved. It is possible that the lack of PA expression may also disrupt this stoichiometry in the newly formed RNP complexes that are formed in the virus-infected cells, and that the correct stoichiometry in the H5N2 virus RNP complex in virus-infected cells may be requirement for the nuclear export of the newly formed structures. In this case the reduced expression of other RNP-associated proteins would be expected to exhibit a similar effect. We therefore examined the effect of the expression of the H5N2 virus PB1 and PB2 proteins on the nuclear export of the RNP complexes in the H5N2 virus-infected cells using the custom-made siRNA libraries.

Cells were treated with siGFP and either the siPB1 (siPB1-1 to siPB1-4) or siPB2 (siPB2-1 to siPB2 -4) libraries and the cells infected with H5N2 virus using a moi of 5 (Fig. 5). At 24 hpi the PB1 mRNA levels (Fig, 5A(i)) and PB2 mRNA levels (Fig, 5A(ii)) were compared with that in non-treated cells as described above. Similar PB1 and PB2 mRNA levels were detected in non-treated and siGFP-treated infected cells in these assays consistent with no inhibitory effect of the siGFP on virus gene expression as indicated above. While cells treated with siPB1-1 and siPB1-3 showed a 70 to 80% reduction in PB1 mRNA levels, the treatment with siPB2-1 to -4 exhibited a 60% reduction in PB1 mRNA levels. In cells treated with siPB1-2 similar PB1 mRNA levels to that recorded in siGFP-treated cells was noted, suggesting no change in PB1 protein expression following siPB1-2 treatment. In contrast, while all the individual siRNA species in the siPB2 library exhibited reduced PB2 mRNA levels, the greatest reduction PB2 mRNA levels was observed in siPB2-1 and siPB2-2-treated cells which showed an 80% reduction in PB2 mRNA levels. We treated cells with each siRNA within the siPB1 library (Fig. 5B(i)) and siPB2 library (Fig. 5B(ii)) together with siGFP (Fig. 5B(iii) and infected the cells with H5N2 virus using a moi of 5. At 24 hpi the cell monolayers were stained using anti-NP and imaged using IF microscopy. While anti-NP staining was detected in cells treated with all siRNA species, the anti-NP staining again appeared to be localized to the nucleus of the infected cells treated with siPB1-1 to siPB1-3, and in infected cells treated with all siPB2 species. This was consistent with the impaired nuclear export of the newly formed RNP complexes as described above. In contrast, in cells treated with siPB1-2, siPB1-4, and siGFP the anti-NP staining appeared to be more diffuse across the entire cells and was consistent with nuclear export of the newly formed RNP complexes. These data again indicated a correlation between the magnitude of the reduction in the PB1 and PB2 mRNA levels and the impaired nuclear export of the RNP complexes in the H5N2 virus-infected cells.

**Figure 5.**
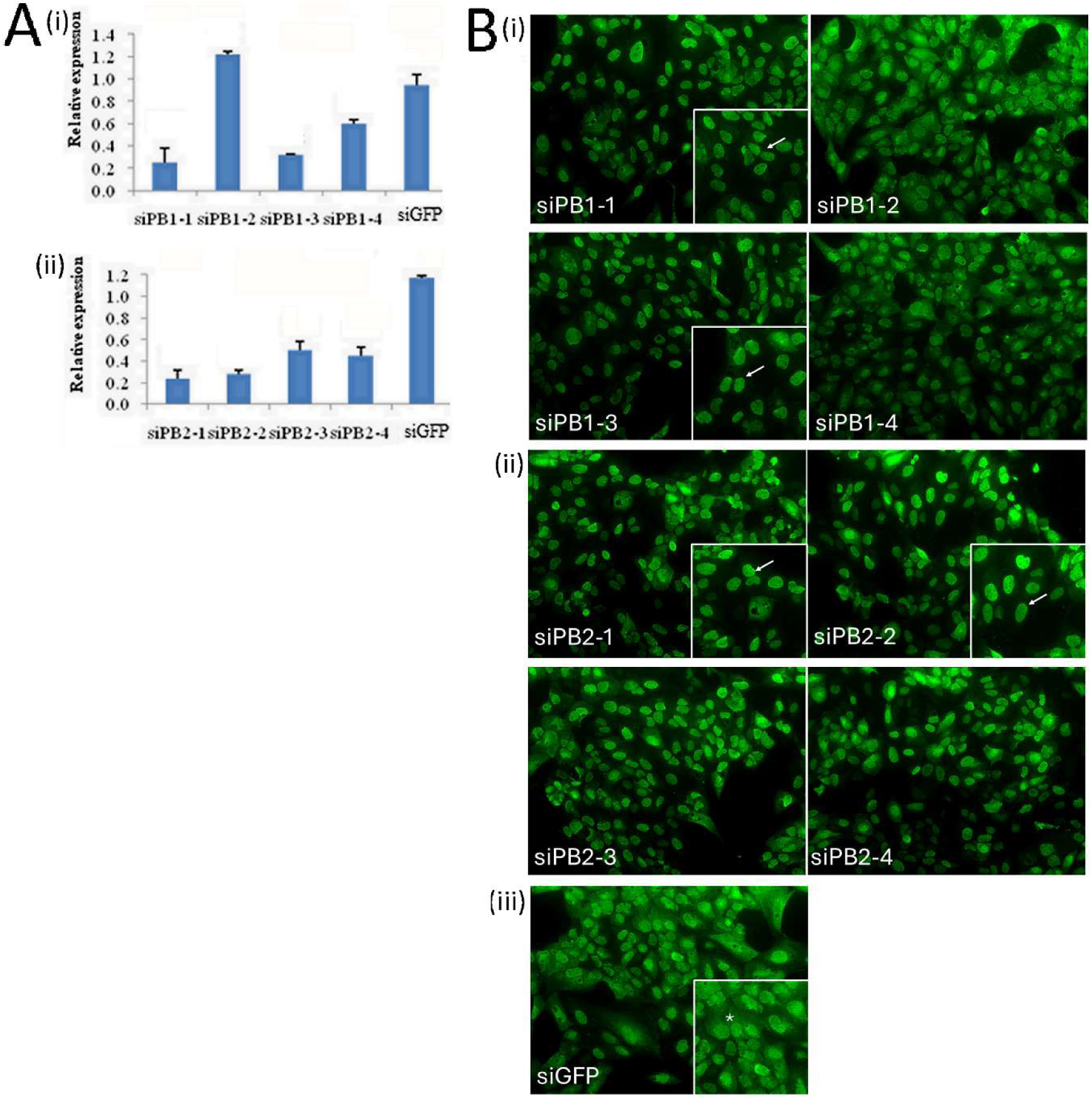
Silencing of the H5N2 virus PB1 and PB2 proteins in H5N2 virus-infected MDCK cells. MDCK cells were treated with either **(A)** (i) siGFP or siPB1-1, siPB1-2, siPB1-3 or siPB1-4 and (ii) siGFP or siPB2-1, siPB2-2, siPB2-3 or siPB2-4 as indicated and infected with the H5N2 virus using the multiplicity of infection of 5. The mRNA coding for the corresponding P(i) B1 mRNA and (ii) PB2 mRNA was measured by qPCR and the relative levels of each is shown with respect to the corresponding mRNA levels in untreated cells is shown. Each reading was performed in triplicate, and the average readings are shown. **(B)** MDCK cells were treated with either each of the (i) siPB1 species, (ii) PB2 species and (iii) siGFP as indicated and infected with the H5N2 using the multiplicity of infection of 5. The cells were stained using anti-NP and imaged by fluorescent microscopy (objective x 40 magnification). The insets in each plate are enlarged images of the cells in the field of view for each plate to more clearly image the anti-NP staining pattern and the nuclear anti-NP staining pattern (white arrows) is highlighted.

#### 2.2.4 The antiviral siRNA that targets the PA, PB1 and PB2 proteins block H5N2 virus transmission

Cells treated with siGFP and siRNA molecules that targeted the PA, PB1 and PB2 proteins were further examined to confirm their antiviral activity using the multiple cycle H5N2 virus infection model described above. In this analysis we selected the two siRNA molecules from each library that showed the highest reduction in the mRNA levels for each target gene (i.e. PA, PB1 and PB2) in the H5N2 virus-infected cells. Cells were treated with siGFP (Fig. 6A) and siPA-3 and siPA-4 (Fig. 6B(i)), siPB1-1 and siPB1-3 (Fig. 6B(ii)) and siPB2-1 and siPB2-2 (Fig. 6B(iii)) and infected with H5N2 virus using a moi of 0.01. At 48 hpi the cell monolayers were co-stained with anti-NP and EB, and the co-stained monolayers were imaged using IF microscopy. The siGFP treated cells exhibited extensive anti-NP staining across the whole monolayer consistent with efficient virus transmission across the cell monolayer. The reduced numbers of anti-NP-stained cells in the cell monolayers treated with each of the siRNA molecules that target the H5N2 virus genes were consistent with impaired virus transmission.

**Figure 6.**
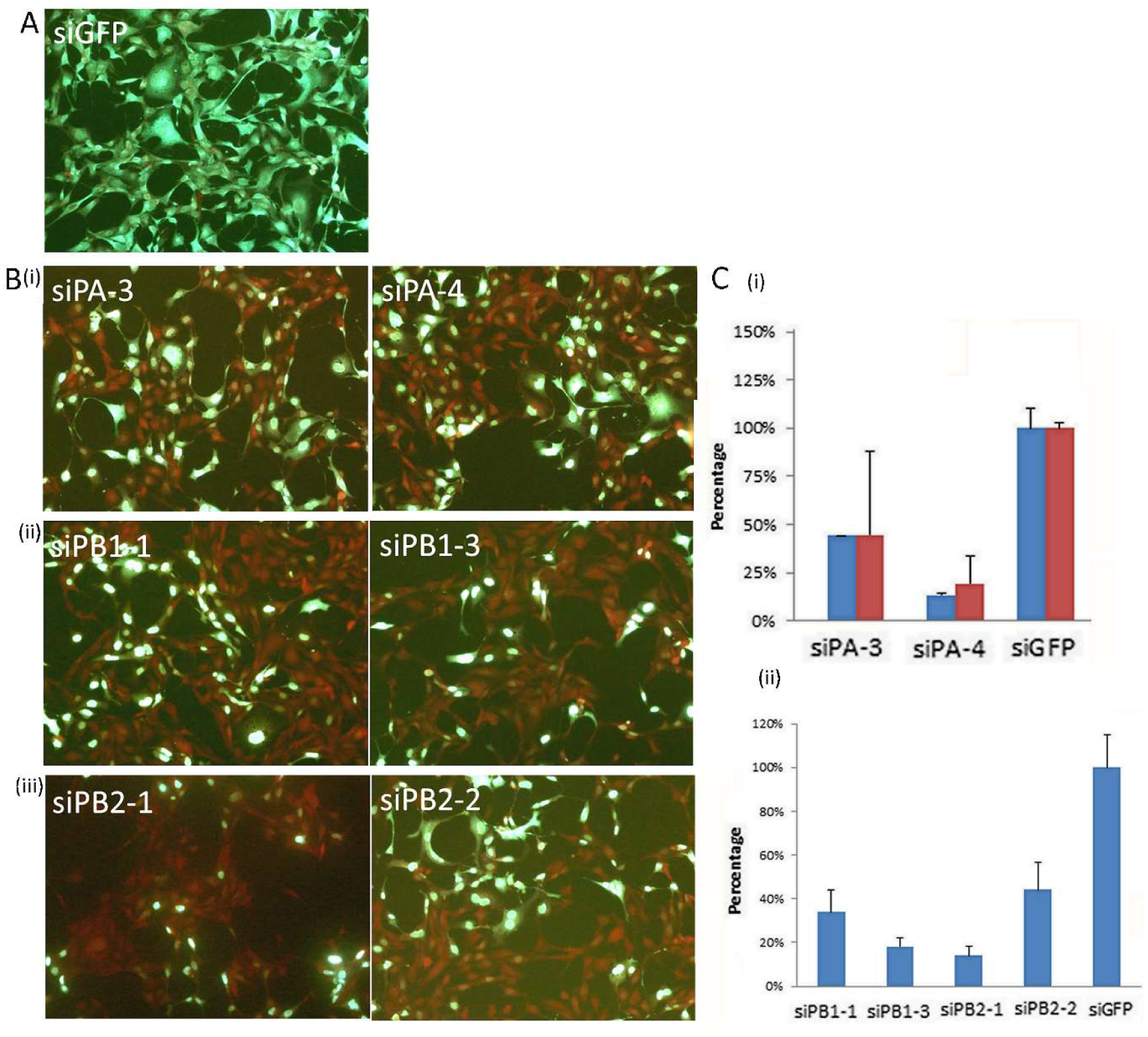
Silencing of the H5N2 virus PA, PB1 and PB2 proteins in H5N2-virus infected MDCK cells blocks virus transmission. MDCK cell monolayers were treated with **(A)** siGFP, **(B) (i)** siPA-3 and siPA-4, **(ii)** siPB1-1 and siPB1-3, and **(iii)** siPB2-1 and siPB2-2 and infected with the H5N2 virus using the multiplicity of infection of 0.01. After 48 hrs post-infection the cells were co-stained using anti-NP (green) and the cell stain Evans blue (red) and imaged using immunofluorescence microscopy (objective x 20 magnification). The Evans blue was used to visualise all the cells in the field of view. **(C)** MDCK cell monolayers were infected with the H5N2 virus using the multiplicity of infection of 0.1. **(i)** Cells were treated with siGFP, siPA-3 and siPA-4 and after 36 hrs post-infection (blue bar) and 48 hrs post-infection (brown bar) the virus infectivity in the tissue culture medium was measured using plaque assay. **(ii)** Cells were treated with siGFP, siPB1-1, siPB1-3, siPB2-1 and siPB2-2 and at 48 hrs post-infection the virus infectivity in the tissue culture medium was measured using plaque assay. In all cases the percentage change relative to that in untreated cells infected with the H5N2 virus is shown and each measurement is an average of three readings.

The virus infectivity that was recovered from cells that were treated with siGFP and siPA-3 and siPA-4 (Fig. 6C(i)) and either siPB1-1 and siPB1-3, or siPB2-1 and siPB2-2 (Fig. 6C(ii)) were measured and compared with that of non-treated cells. A 60 to 90% reduction in the virus infectivity in the cells treated with each of the siRNA molecules that target the virus genes was observed, while no reduction in virus infectivity from the siGFP-treated cells was recorded. These data obtained using the multiple cycle infection model confirmed the antiviral activity of the siRNA molecules that target the virus genes and confirmed that these siRNA molecules block virus transmission.

#### 2.2.5 The siRNA targeting the H5N2 NP and polymerase proteins also inhibit the activity of the H5N2 polymerase complex

In a final analysis we examined the effect of these siRNA molecules on the virus polymerase (pol) activity using a H5N2 virus mini-replicon assay in human HEK 293T cells (Fig. 7). The minireplicon system was constructed by inserting the H5N2 virus NP gene and the genes for the individual H5N2 virus P proteins into the mammalian expression vector pCAGGS to generate the four expression plasmids pCAGGS-H5N2 NP, pCAGGS-H5N2PA, pCAGGS-H5N2PB-1 and pCAGGS-H5N2PB2. The expression of the H5N2 NP and the individual P proteins in human HEK 293T cells was confirmed using appropriate antibodies. Immunoblotting analysis revealed expressed virus proteins of the expected sizes (SFig. 2A-D) and IF microscopy that showed high levels of transfection with each plasmid in these cells (SFig. 2E). Cells were co-transfected with the four expression plasmids and pPolL1-Luc and the pol activity of the H5N2 polymerase complex was assessed in cells expressing the H5N2 virus polymerase complex using luciferase expression as a reporter assay to indicate pol activity. This analysis was performed in non-treated cells and in cells that were treated with either siGFP and. either siNP-1, siNP-2, siNP-3 or siNP-4 (Fig. 7A) and siPA-3, siPB1-3 and siPB2-1 (Fig. 7B). The measurements were made at 3 days post-transfection which was the peak activity in non-treated cells expressing the recombinant H5N2 virus polymerase complex (SFig. 3).

**Figure 7.**
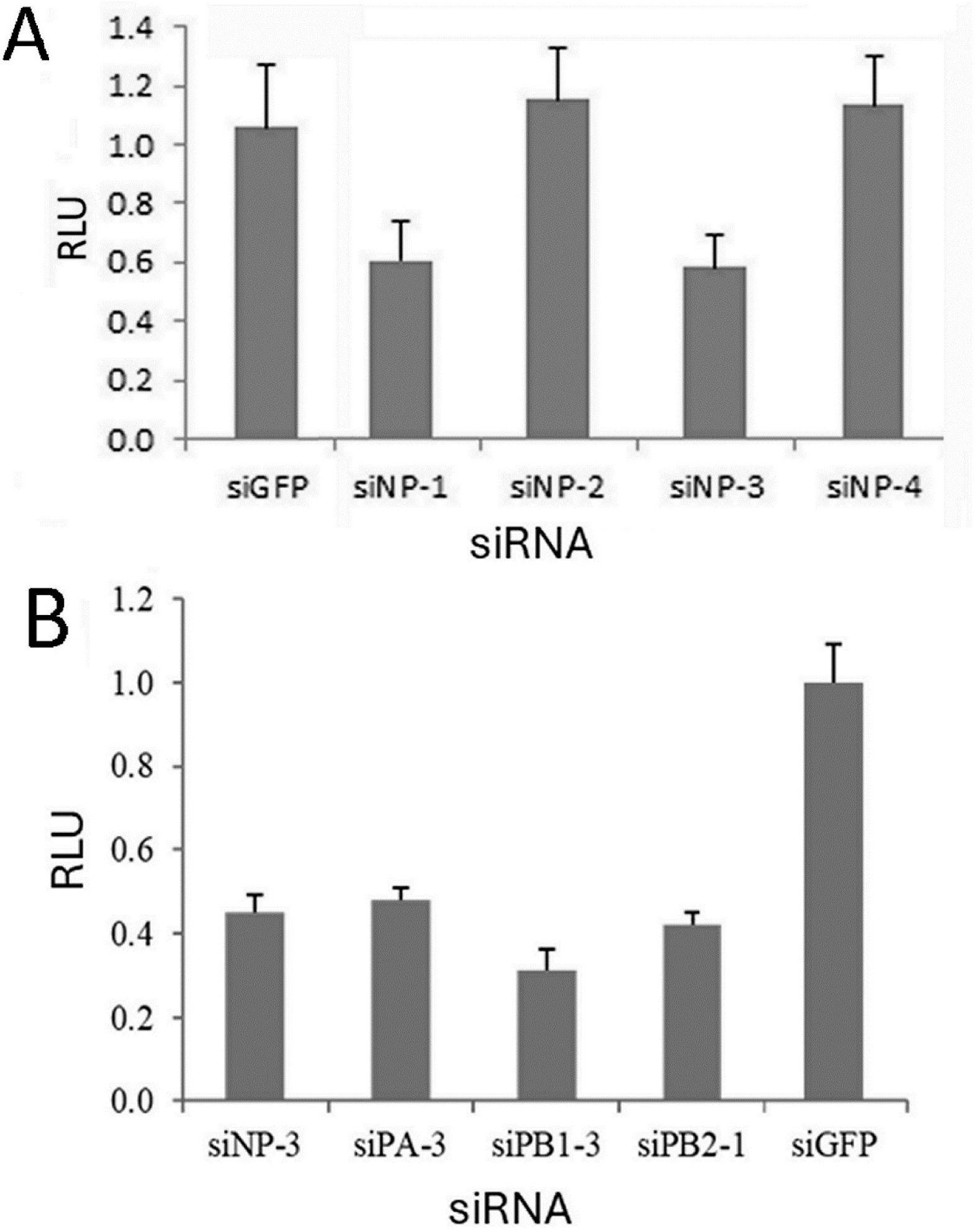
The antiviral siRNA species inhibit activity of the H5N2 polymerase complex. The polymerase activity of the recombinant H5N2 polymerase complex was measured in HEK 293Tcells treated with **(A) (i)** siGFP and each of the H5N2-specific NP siRNA species as indicated and **(B)** siGFP and selected H5N2-specific NP siRNA species that target the NP (NP-3), PA (siPA-3), PB1 (siPB1-3) and PB2 (siPB2-1) protein expression. In each case the polymerase activity (luciferase units) relative to that in untreated cells expressing the respective polymerase complex in shown (RLU). Each measurement is an average of three readings.

While similar levels of H5N2 virus polymerase activity were recorded in non-treated cells and in cells treated with either siGFP, siNP-2 and siNP-4, a 40% reduction in the H5N2 virus pol activity was recorded in the cells treated with siNP-1 and siNP-3 (Fig. 7A). These observations in the pol activity assay correlated with the data obtained in H5N2 virus-infected cells that indicated siNP-1 and siNP-3 exhibited the greatest reduction in NP mRNA levels. These data provided additional evidence that the respective siNP species target the NP for each virus, and as expected confirmed that the presence of the NP is required for the activity of newly formed influenza virus polymerase complexes in cells.

Since siPA-3, siPB1-3 and siPB2-1 exhibited greatest anti-viral activity we also used these as representative siRNA molecules from the PA, PB1 and PB2 siRNA library to examine their effects on the polymerase activity in cells expressing the H5N2 virus polymerase complex (Fig. 7B). In this analysis the effects of siPA-3, siPB1-3 and siPB2-1 on the pol activity were compared in parallel with that in cells treated with siNP-3 and siGFP. While the siGFP-treated cells exhibited no inhibitory effect on the H5N2 virus pol activity, in the cells treated with siNP-3 an approximate 60% reduction in pol activity was recorded. The cells treated with siPA-3 exhibited an approximate 50% in polymerase activity, while in the cells treated with siPB1-3 and siPB2-1 a 60-70% reduction in polymerase activity was recorded. This general reduction in polymerase activity with siPA-3, siPB1-3 and siPB2-1 was similar to the reduction in pol activity exhibited in the siNP-3-treated cells. This analysis confirmed the requirement of the PA, PB1 and PB2 proteins in establishing a functional polymerase complex and correlated with the antiviral effects of these molecules in the H5N2 virus-infected cells. The inhibition in the pol activity in the cells treated with the respective the siPA, siPB1 and siPB2 also provided additional evidence that the anti-NP staining that was observed in the nucleus of H5N2 virus-infected cells treated with the siPA, siPB1 and siPB2 resulted from the transcriptional activity of the fully functional a pre-formed polymerase complex that arises from the input H5N2 virus (i.e. from the inoculum).

In this report we have noted that while effective siRNA species were identified in the SMARTpool libraries from each target virus gene, a minor proportion of the siRNA species generated showed no antiviral effect, which correlated with moderate/no effect on reducing the virus pol activity. The *invitro* polymerase assay may therefore represent a general strategy with which to screen the SMARTpool libraries that target the virus polymerase genes prior to engaging in live virus work, which may be an important consideration when high containment facilities are required for the virus work.

## 3. Conclusion

We have previously described the biological properties of a low passaged tissue culture grown avian virus A/Duck/Malaysia/02/2001 (H9N2) that was isolated from live poultry during routine surveillance [19]. The H9N2 virus could be readily propagated using embryonated eggs and in avian cell culture (e.g. CEF cell culture), and although the H9N2 virus could efficiently infect mammalian cell lines (e.g. MDCK cells) it could not be propagated using these cell types [18]. An inhibition in the nuclear export of the newly formed RNP complexes in mammalian cells infected with the H9N2 virus was noted that did not occur in avian cells. This suggested that the impaired nuclear export formed the basis for the restricted transmission and propagation of the H9N2 virus in these non-avian cells [20]. This impaired nuclear export of the H9N2 virus RNP in these cells correlated with the severe reduced expression of the PA protein in non-avian cells such as the A549 and MDCK cells. Although the mechanism behind the reduced expression of the PA protein in non-avian cells is currently unclear, it appeared to an intrinsic property of the H9N2 virus PA protein. In this study we tested the prediction that the reduced expression of the H5N2 PA protein would also lead to a phenotype in H5N2 virus-infected MDCK cells that was similar to that previously observed in H9N2 virus-infected MDCK cells. Although the H5N2 virus described above can be propagated in the MDCK cells line; by artificially silencing the expression of the H5N2 PA protein using a siRNA strategy we have replicated this same phenomenon that was observed in H9N2 virus-infected MDCK cells i.e. nuclear retention of the newly formed H5N2 virus RNP complexes. Our current study provides evidence to support our earlier suggestion that the reduced expression of the PA protein in mammalian cells infected with the circulating A/Duck/Malaysia/02/2001 (H9N2) leads to impaired nuclear export of the H9N2 virus RNP complex, and is a factor that restricts virus transmission in some H9N2 virus strains that circulate in the natural environment. This may reflect a failure of the H9N2 virus to adapt to the non-avian cell background and that leads to the inhibitory expression of the H9N2 virus PA protein in non-avian cells. This finding warrants a better understanding of mechanism involved to determine if this is a failure to interact with specific cellar factors that are required for PA protein expression e.g. the interaction of the virus mRNA with host factors in the translational machinery. An understanding of the molecular basis for this attenuation of the H9N2 virus in mammalian cells could facilitate the development of attenuated influenza virus vaccines that can be used in non-avian species by using reverse genetics to introduce specific sequence changes into the PA gene of the influenza virus vaccine candidates. These engineered viruses would be expected to be readily propagated using standard egg culture but would be attenuated in non-avian hosts.

Artificially silencing the expression of the H5N2 PA protein in H5N2 virus-infected cells using siPA lead to the nuclear retention of the newly formed H5N2 virus RNP complexes in virus-infected cells and a block in virus transmission. However, a similar impaired nuclear export of the newly formed H5N2 virus RNP complexes were also observed following silencing of the expression of the H5N2 PB1 and PB2 proteins, which suggests that equimolar expression of all the virus P proteins may be required for nuclear export of the newly formed RNP complexes. It is currently unclear if this is related to specific properties associated with the individual virus P proteins vis-a-via nuclear export of the RNP complex, or if the interaction with the individual P proteins with the RNP complex is a structural requirement that allows the adoption of the correct protein complex conformation to enable nuclear export of the RNP complex. Although these possibilities will require further investigation, our current study has highlighted the potential use of siRNA strategies that target the virus polymerase as an antiviral strategy to control avian influenza virus infection in the airway of non-avian hosts. This siRNA approach would only require sequence information relating to the virus in question, and its application as an antiviral strategy should be facilitated by the recent development of carrier systems for the delivery of RNA-based vaccines.

## Supporting information

extra data

## Acknowledgements

We acknowledge Dawn S. Yeo and Myint Zu Myaing for technical assistance. We thank the Singapore Ministry of Education (MOE) for the award of a PhD scholarship to Richard Sutejo. We thank the Agri-Food & Veterinary Authority of Singapore (Lim Chee Wee and Yueh Nuo Lin) for providing the H5N2 avian virus. This work was supported by grants from Défense Science Technology Agency Singapore (BHT), and the National Medical Research Council of Singapore (RJS) and MOE (RJS).

## Supp Figure legends

**SFig. 1 Monitoring the anti-NP staining in a time course of infection following H5N2 virus infection.** MDCK cells were **(A)** mock-infected and **(B-D)** infected with the H5N2 virus. At **(B)** 8 hrs post infection, **(C)** 10 hrs post-infection and **(D)** 16 hrs post-infection the cells were co-stained with anti-NP and Evans blue (EB). The co-stained cells were imaged using immunofluorescence microscopy ((objective x 20 magnification). The anti-NP nucleus-staining pattern (white arrow) and cytoplasmic staining patterns (*) are highlighted.

**SFig. 2 Expression of the recombinant H5N2 virus NP and P proteins in HEK 293T cells.** Cells were mock-transfected with pCAGGS (M) and transfected (T) with (**A)** pCAGGS/NP, **(B)** pCAGGS/PA, **(C)** pCAGGS/PB1 and **(D)** pCAGGS/PB2 and at 36 hrs post-transfection the cells were extracted in boiling mix and examined by immunoblotting with the appropriate antibody. Protein species corresponding to the corresponding full-length protein are indicated. **(E)** is the loading control using anti-actin. **(F)** Cells were transfected with pCAGGS only and (i) pCAGGS/NP, (ii) pCAGGS/PA, (iii) pCAGGS/PB1 and (iv) pCAGGS/PB2 and at 36 hrs post-transfection the cells were stained using the appropriate antibody and examined by immunofluorescence microscopy (objective 20x magnification).

**SFig. 3. Polymerase activity in cells expressing the polymerase complex. (A)** The luciferase activity in HEK 293T cells co-transfected with pCAGGS and pPolL1-Luc only (blue column) and with pCAGGS-H5N2 NP, pCAGGS-H5N2 PA, pCAGGS-H5N2 PB-1 and pCAGGS-H5N2 PB2 that expresses the recombinant H5N2 polymerase complex and pPolL1-Luc (purple column) was measured over six days. In each case the polymerase activity (luciferase units) relative to that in non-transfected cells in shown (RLU). Each measurement is an average of three readings. The respective relative luciferase activity in shown (RLU) and each measurement is an average of three readings. **(B)**. HEK 293T cells co-transfected with (i) pCAGGS and pPolL1-Luc only and (ii) recombinant H5N2 polymerase complex and pPolL1-Luc (purple column) and at 48 hrs post-transfection the cells were co-stained using anti-NP and Evans blue and the cells examined by immunofluorescence microscopy (objective 20x magnification).

**STable. 1. Primers used in the qPCR analysis of H5N2 virus infected cells.** The primer pairs used for determining the expression of the H5N2 virus NP, PA, PB1 and PB2 mRNAs in virus infected cells are indicated. The primer pairs used to detect the Canine EF is also shown which was used to normalise the assays. The forward (FW), reverse (RV) and UPL number are indicated for each gene.

**STable. 2. Primers used for subcloning of the H5N2 NP and polymerase genes into the pCAGGS expression vector.** The forward (FW) and reverse (RV) primer pairs are indicated and the restriction enzyme site underlined.

## References

1. Webster RG, Bean WJ, Gorman OT, Chambers TM, Kawaoka Y (1992). Evolution and ecology of influenza A viruses. Microbiol Rev 56: 152–179.Evolution and ecology of influenza A viruses. Microbiol Rev 56: 152–179.

2. Peiris M, Yuen KY, Leung CW, Chan KH, Ip PL, et al. (1999) Human infection with influenza H9N2. Lancet 354: 916–917.

3. Butt KM, Smith GJ, Chen H, Zhang LJ, Leung YH, et al. (2005) Human infection with an avian H9N2 influenza A virus in Hong Kong in 2003. J Clin Microbiol 43: 5760–5767.

4. Ogata T, Yamazaki Y, Okabe N, Nakamura Y, Tashiro M, et al. (2008) Human H5N2 avian influenza infection in Japan and the factors associated with high H5N2-neutralizing antibody titer. J Epidemiol 18: 160–166.

5. Wang M, Fu CX, Zheng BJ (2009) Antibodies against H5 and H9 avian influenza among poultry workers in China. N Engl J Med 360: 2583–2584.

6. Yamazaki Y, Doy M, Okabe N, Yasui Y, Nakashima K, et al. (2009) Serological survey of avian H5N2-subtype influenza virus infections in human populations. Arch Virol 154: 421–427.

7. Boivin, S.; Cusack, S.; Ruigrok, R.W.; Hart, D.J. Influenza A virus polymerase: Structural insights intoreplication and host adaptation mechanisms. J. Biol. Chem. 2010, 285, 28411–28417.

8. Pflug, A.; Guilligay, D.; Reich, S.; Cusack, S. Structure of influenza A polymerase bound to the viral RNA promoter. Nature 2014, 516, 355–360.

9. Reich, S.; Guilligay, D.; Pflug, A.; Malet, H.; Berger, I.; Crépin, T.; Hart, D.; Lunardi, T.; Nanao, M.; Ruigrok, R.W.;, et al. Structural insight into cap-snatching and RNA synthesis by influenza polymerase. Nature 2014, 516, 361–366.

10. Chang, S.; Sun, D.; Liang, H.; Wang, J.; Li, J.; Guo, L.; Wang, X.; Guan, C.; Boruah, B.M.; Yuan, L.;, et al. Cryo-EM structure of influenza virus RNA polymerase complex at 4.3 A resolution. Mol. Cell 2015, 57, 925–935.

11. Coloma, R.; Valpuesta, J.M.; Arranz, R.; Carrascosa, J.L.; Ortín, J.; Martín-Benito, J. The structure of a biologically active influenza virus ribonucleoprotein complex. PLoS Pathog. 2009, 5, e1000491.

12. Schrauwen, E.J.; Fouchier, R.A. Host adaptation and transmission of influenza A viruses in mammals. Emerg. Microbes. Infect. 2014, 3, e9.

13. Manz, B.; Schwemmle, M.; Brunotte, L. Adaptation of avian influenza A virus polymerase in mammals to overcome the host species barrier. J. Virol 2013, 87, 7200–7209.

14. Long, J.S.; Giotis, E.S.; Moncorge, O.; Frise, R.; Mistry, B.; James, J.; Morisson, M.; Iqbal, M.; Vignal, A.; Skinner, M.A.;, et al. Species di_erence in ANP32A underlies influenza A virus polymerase host restriction. Nature 2016, 529, 101–104.

15. Gabriel, G.; Klingel, K.; Otte, A.; Thiele, S.; Hudjetz, B.; Arman-Kalcek, G.; Sauter, M.; Shmidt, T.; Rother, F.; Baumgarte, S.;, et al. Di_erential use of importin-alpha isoforms governs cell tropism and host adaptation of influenza virus. Nat. Commun. 2011, 2, 156.

16. Biswas, S.K.; Boutz, P.L.; Nayak, D.P. Influenza virus nucleoprotein interacts with influenza virus polymerase proteins. J. Virol. 1998, 72, 5493–5501.

17. Paterson, D.; Fodor, E. Emerging roles for the influenza A virus nuclear export protein (NEP). PLoS Pathog 2012, 8, e1003019.

18. Kumar, S., Yeo, D. Muralidharan, N. Lai, S.K., Tong, C. Tan, BH., and Sugrue, R.J. Impaired Nuclear Export of the Ribonucleoprotein Complex and Virus-Induced Cytotoxicity Combine to Restrict Propagation of the A/Duck/Malaysia/02/2001 (H9N2) Virus in Human Airway. Cells, 2020 3;9(2):355.

19. Yeo, D.S.; Ng, S.H.; Liaw, C.W.; Ng, L.M.; Wee, E.J.; Lim, E.A.; Seah, S.L.; Wong, W.K.; Lim, C.W.; Sugrue, R.J. Tan, BH. Molecular characterization of low pathogenic avian influenza viruses, isolated from food products imported into Singapore. Vet. Microbiol. 2009, 138, 304–317.

20. Sutejo, R.; Yeo, D.S.; Myaing, M.Z.; Hui, C.; Xia, J.; Ko, D.; Cheung, P.C.; Tan, B.H.; Sugrue, R.J. Activation of type I and III interferon signalling pathways occurs in lung epithelial cells infected with low pathogenic avian influenza viruses. PLoS ONE 2012, 7, e33732.

21. Myaing, M.; Jumat, R.; Huong, T.; Tan, B.H.; Sugrue, R.J. Truncated forms of the PA protein containing onlythe C-terminal domains are associated with the RNP complex within H1N1 influenza virus particles. J. Gen.Virol 2017, 98, 906–921.

22. Ge, Q., Filip, L., Bai, A., Nguyen, T. Eisen, H.N and Chen, J. Inhibition of influenza virus production in virus-infected mice by RNA interference Proc Natl Acad Sci U S A (2004). 101(23):8676–81.

23. Spackman E, Senne DA, Myers TJ, Bulaga LL, Garber LP, et al. Development of a real-time reverse transcriptase PCR assay for type A influenza virus and the avian H5 and H7 hemagglutinin subtypes. J Clin Microbiol. 2002;40:3256–3260. doi: 10.1128/JCM.40.9.3256-3260.2002

