## Supplementary material for "Evaluating the use of siRNA to silence the expression of the H5N2 virus polymerase genes as strategy to block the transmission of the avian H5N2 virus in mammalian cells": extra data

S Figure 1

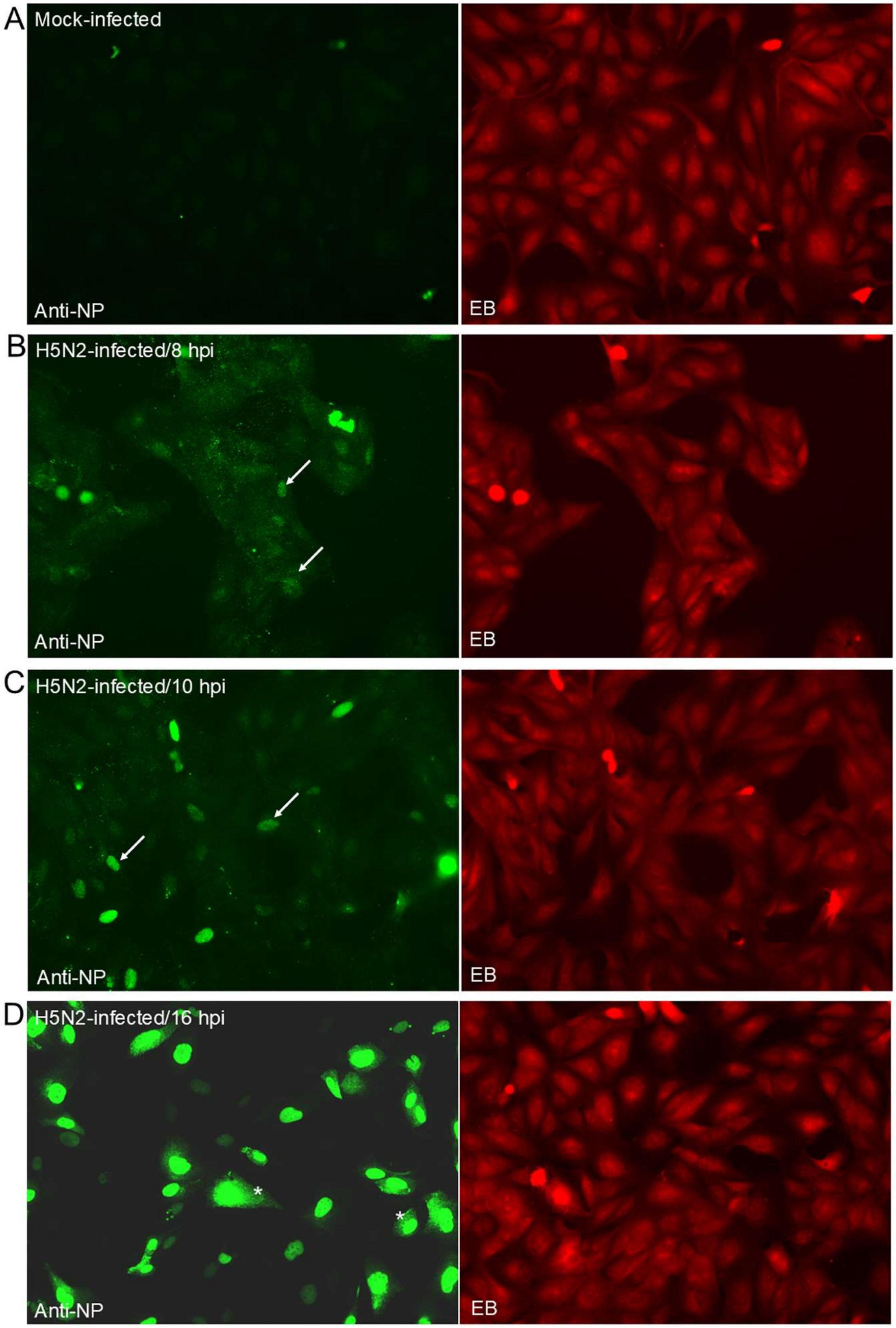
A
Mock-infected
Anti-NP
EB
B
H5N2-infected/8 hpi
Anti-NP
EB
C
H5N2-infected/10 hpi
Anti-NP
EB
D
H5N2-infected/16 hpi
*
*
Anti-NP
EB

S Figure 2

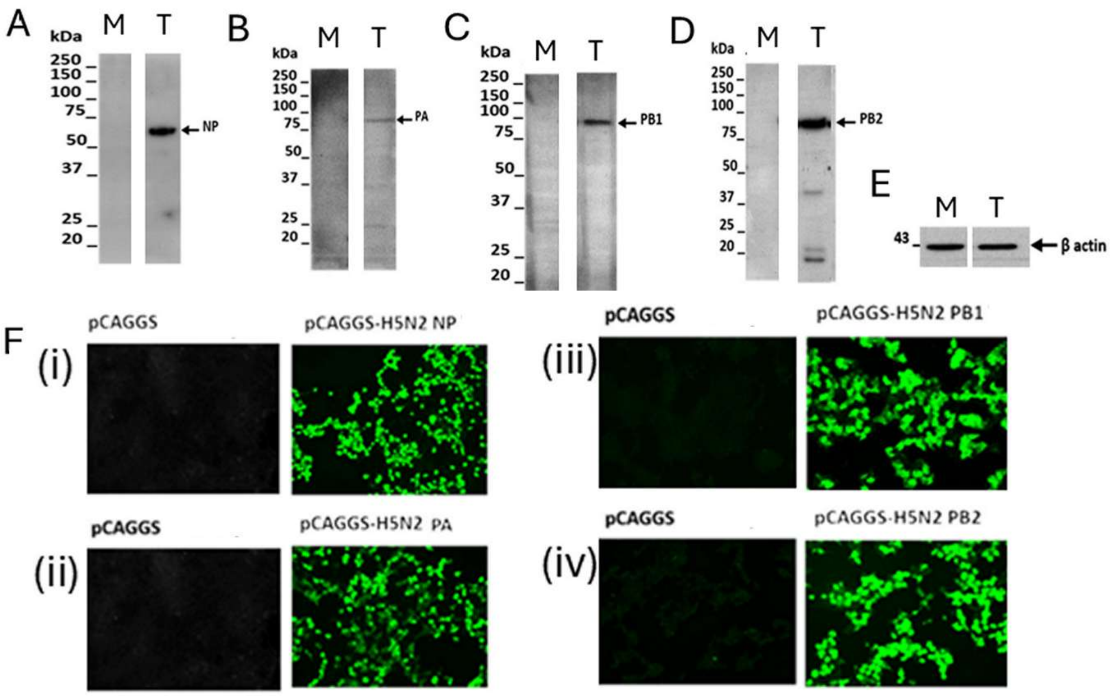

A
kDa
M
T
250
150
100
75
50
37
25
20
NP
B
PA
C
PB1
D
PB2
E
43
β actin
F
(i)
(ii)
(iii)
(iv)
pCAGGS
pCAGGS-H5N2 NP
pCAGGS-H5N2 PA
pCAGGS-H5N2 PB1
pCAGGS-H5N2 PB2

S Figure 3

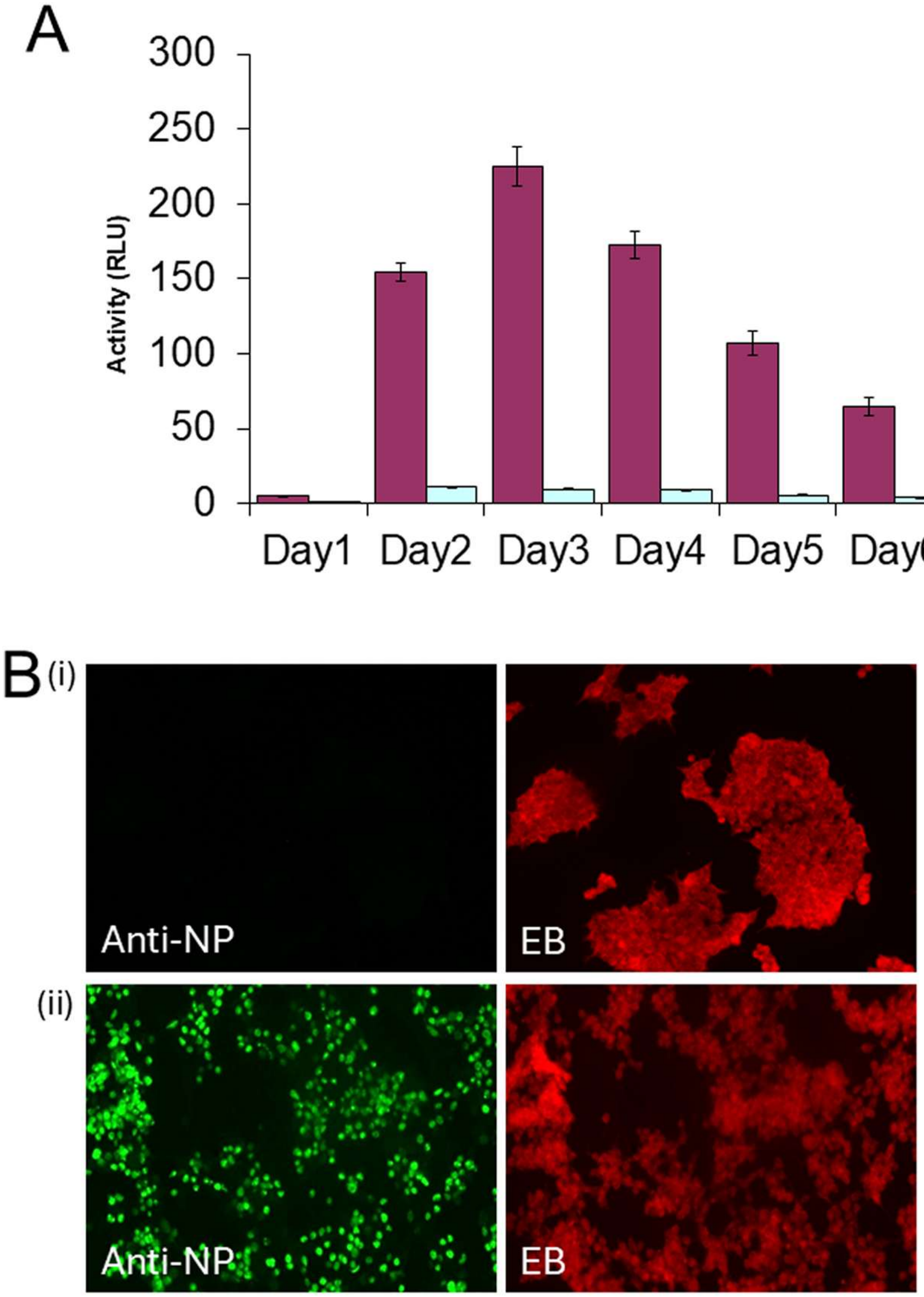

A
300
250
200
150
100
50
0
Activity (RLU)
Day1
Day2
Day3
Day4
Day5
Day6
B
(i)
Anti-NP
EB
(ii)
Anti-NP
EB

S table 1

| Virus | Gene | Sequence (5'-3') | Probe Sequence (UPL Probe #) |
|---|---|---|---|
| H5N2 | NP FW | TGCTTCAAATGAGAACATGGA | CTCCAGCA (#67) |
| | NP RV | GCCCAATATCTGCTTCTTAGTTCA | |
| | PA FW | GGTATAAACCCAAATTACCTCCTG | GGAAGCAG (#38) |
| | PA RV | AATGTCTTGGAGTTCTGCCAGT | |
| | PB1 FW | ATACAGGAGGCCGGTTGG | TGGTGGAG (#22) |
| | PB1 RV | ATTCGGGCCCTAGACACC | |
| | PB2 FW | AGAGCAACGGCCATTCTAAG | GCAACCAG (#164) |
| | PB2 RV | TTCAGCAATTGATTGTTCGTCT | |

| Host | Gene | Sequence (5'-3') | Probe Sequence (UPL Probe #) |
|---|---|---|---|
| Canine | EF FW | GCTGGAAGATGGTCCCAAG | GGATGCTG (#89) |
| | EF RV | TGCCAGGAACCATATCAACA | |

S table 2

| H5N2/F118 virus genes | Primers | Sequence (5'-3') |
|---|---|---|
| NP | NP-EcoRI-FW | CCGGAATTCCGGAGCAAAAGCAGGGTA |
| | NP-KpnI-RV | CGGGGTACCCCGAGTAGAAACAAGGGTATTTTT |
| PA | PA-KpnI-FW | CGGGGTACCCCGAGCGAAAGCAGGTAC |
| | PA-KpnI -RV | CGGGGTACCCCGAGTAGAAACAAGGTACTT |
| PB1 | PB1-KpnI-FW | CGGGGTACCCCGAGCGAAAGCAGGCA |
| | PB1-XhoI-RV | CCGCTCGAGCGGAGTAGAAACAAGGCATTT |
| PB2 | PB2-KpnI-FW | CGGGGTACCCCG AGCGAAAGCAGGTC |
| | PB2-XhoI-RV | CCGCTCGAGCGGAGTAGAAACAAGGTCGTTT |
